# Pedigree Painter (PePa): a tool for the visualization of genetic inheritance in chromosomal context

**DOI:** 10.1101/2024.12.18.629215

**Authors:** Andrea Pozzi

## Abstract

**Background:** Data visualization is increasingly important in genomics, enabling researchers to uncover inheritance and recombination patterns across generations. While most existing tools focus on ancestry prediction, they lack functionality for analyzing known ancestries in controlled settings, such as determining parental contributions to offspring genomes. To address this gap, I developed *pepa*, a lightweight, modular tool that visualizes and quantifies genomic inheritance, designed for beginner and advanced users.

**Results:** *pepa* is a program for processing VCF files, assigning ancestries to SNPs, and clustering them into biologically meaningful regions. It generates human-readable comparison tables and visualizes inheritance patterns with chromosome paintings through R. Tested on fission yeast, *pepa* revealed non-uniform recombination patterns, with chromosomes largely inherited from one parent and seemingly random recombination. Quantitative analyses showed differences in parental contributions at the nucleotide and gene levels, with some offspring inheriting similar percentages from parents. However, the painted chromosomes revealed that even offspring with similar percentages from one parent rarely inherit the same genomic region, highlighting the importance of this tool in drawing biologically meaningful insights.

**Conclusion:** *pepa* provides an accessible and powerful solution for analyzing genomic inheritance, bridging experimental and computational biology. Its modular design and minimal dependencies allow adaptation to diverse organisms, facilitating intuitive visualization and quantitative insights into recombination dynamics.

## Introduction

The importance of data visualization is rising in biology, as we accumulate an increasing amount of genomic data [1]. Visualizing data is a cornerstone of genomics [2], enabling researchers to uncover patterns that would otherwise remain unknown. This is particularly important for understanding genome recombination across generations, a topic important across multiple fields, from experimental biology to human evolution [3, 4]. Recombination shapes genomes by mixing parental contributions, but the effects observed over a long time (ancestry) often differ from those happening over a short time, for example, from parents to offspring (pedigree). While the pedigree of an organism is influenced by its ancestry, sometimes the interest focuses on simply understanding from which parent-specific parts of the genome can be inherited. One such example is yeast research, where new hybrids are generated multiple times trying to select specific phenotypes of interest using parental strains with phenotypes of interest.

There are many ancestry prediction and visualization tools, such as ChromPlot [5] or human-specific commercial tools like Chromosome Painter (AncestryDNA®). Most tools focus on ancestry prediction and thus lack versatility in showcasing known ancestries [6, 7], such as parental contributions to offspring in a controlled setting. This is best exemplified by the software STRUCTURE [8], used by most researchers in population genetics, and with several tools to plot its predicted ancestries [7]. However, similar programs are not helpful when ancestries are known (e.g., known parents) and the goal is to measure the contribution of each parent to an offspring genome. Such types of experiments are relatively common. For example, researchers working on the evolution of cichlids often resort to ‘common garden experiments’ where the offspring of specific individuals is under study. Another example is the studies of population genetics performed on yeast populations, in which strains are crossed to investigate how the offspring inherit specific phenotypes. This tool aims to fill this gap and provide an accessible solution for analyzing how much of the genome is inherited from each parent, and which specific genomic regions, or genes, are passed down.

### Implementation

#### Pipeline Design

*Pepa* is a versatile and accessible tool for visualizing genomic inheritance, recombination patterns, and parental contributions (**Fig.1**). Its design prioritizes ease of use and customization, allowing both beginners and advanced users to use this tool to analyze their data. The pipeline is developed using Bash, Python, and R. Bash serves as the backbone, connecting various Python and R scripts, making it easy to add extra tools to the pipeline. Bash was used instead of Python as most biologists have at least basic knowledge of Bash. The graphics are computed using R, a common language for graphics in biology, with use spanning from transcriptomics [9, 10] to population genetics [5, 11]. To make this tool useful to more advanced users, the output tables are saved and optimized for R packages such as ggplot2, allowing users to easily customize their plots with only knowledge of R. The pipeline is lightweight and requires minimal installation. The core tool (*pepa-base*) only requires Python 3, without extra modules or installation. The extended pipeline, which includes data visualization (*pepa-paint*), relies on three common R packages to generate graphics.

**Figure 1.**
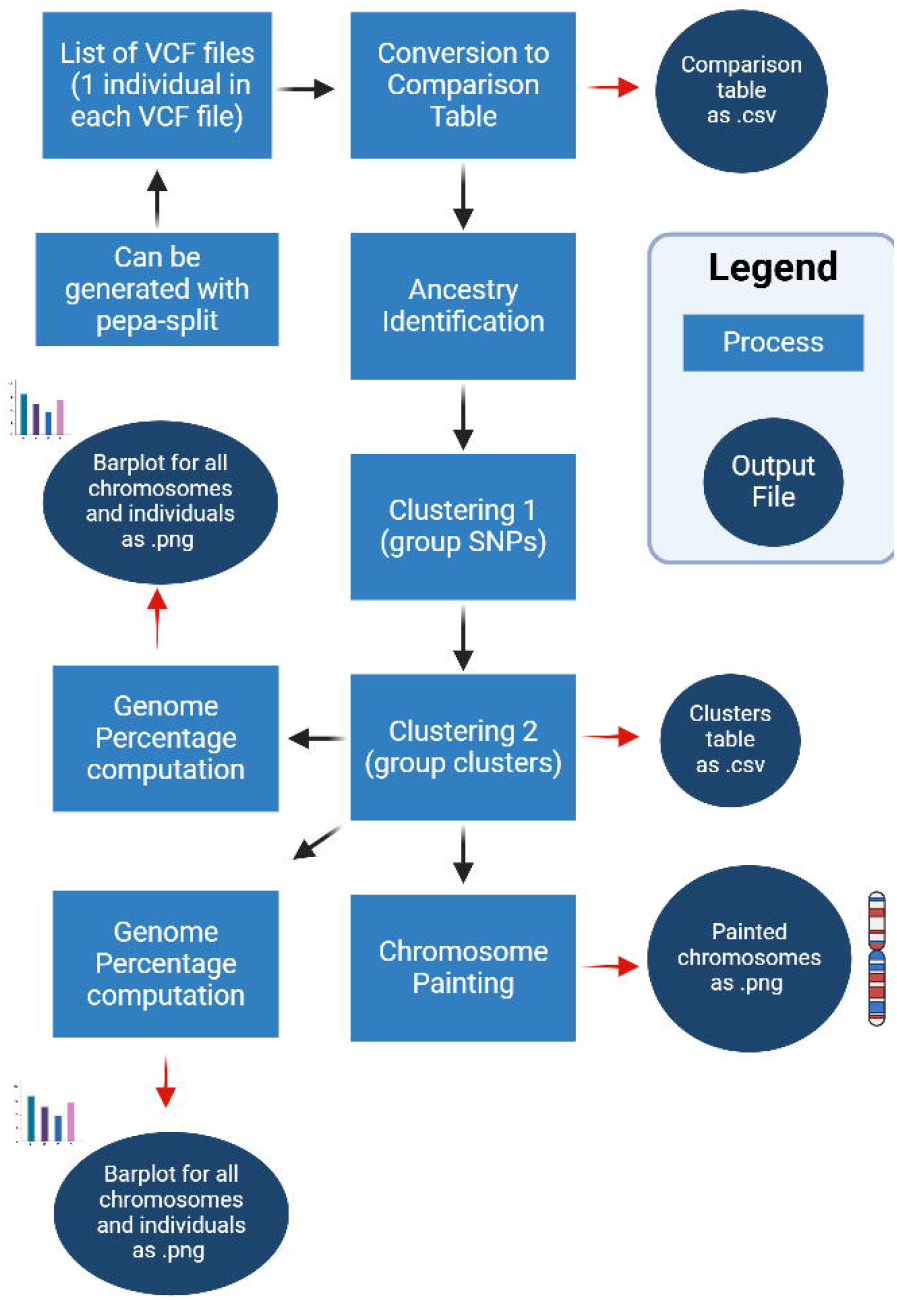
Overview of the *pepa* workflow. *pepa* processes VCF files to generate comparison tables, identify ancestries, and cluster SNPs into biologically meaningful regions. The pipeline produces both plots, such as painted chromosomes, and tables optimized for ggplo2 for custom visualization.

#### Comparison Tables and Ancestry Identification

The first script of *pepa* generates comparison tables summarizing genetic similarities and differences in a human-readable format. This functionality mirrors that of tools like ***vcftoolz*** *[12]*, but with the advantage of requiring no additional installations or dependencies. Each SNP in this table is then assigned an ancestry, from either of the parents. These parents are indicated in the command line, and they can be actual parents (eg. mother and father), or, when working on hybrid species, they can be two representative genomes from those species. The latter option is valuable when analyzing individuals from a hybrid zone, such as seen with crows in Europe [13]. The table generated is then used to compare SNPs between parents and offspring, keeping only SNPs that are presented in either of the parents and discarding all the others. It then assigns the ancestry based on which parent is matched for that specific SNP. The script loads only part of the SNPs into the memory at each time, making the process slower but allowing analyses on machines with low memory. However, thanks to the data structure and multithreading, millions of SNPs across dozens of samples can be converted in less than 10 minutes on a home desktop.

#### Clustering

*Pepa* includes clustering functionality to group SNPs with similar ancestry. While comparison tables show the ancestry of individual SNPs, it is computationally intensive to plot figures with millions of data points, thus, *pepa* includes two clustering algorithms to group close SNPs. In the first clustering algorithm, continuous SNPs with identical ancestry are clustered. Users can define the minimum length of a cluster to be annotated, allowing the exclusion of biologically insignificant clusters caused by erroneous SNP calls (eg. 10nt cluster). In the second clustering algorithm, *pepa* combines non-continuous clusters, eliminating small clusters of different ancestry between two regions with the same ancestry. The size of the clusters to ignore is decided by the user. This process eliminates small clusters (eg. 100nt) between long clusters (eg. Parent1) of the opposite ancestry (eg. Parent2) as these are unlikely to be recombination events. For example, if two clusters of 10kb each belong to Parent1 but a small 100nt cluster in the middle belongs to Parent2, this algorithm assumes that this Parent2 cluster is not real and eliminates it, creating a 20kb Parent1 cluster. In many species, it is unlikely to find many recombination events of a few hundred nucleotides, and this method helps remove these false small recombination events.

#### Chromosome Painting

Using R, *pepa-paint* generates visual representations of chromosomes, highlighting regions inherited from each parent. The code uses three R packages, *dplyr, ggplot2*, and *patchwork* to make the graphics of each individual and chromosome to then plot them into a single plot. The code is optimized to plot genomes with a few chromosomes (<10) across medium size cohorts (<100), however, the native code generates understandable plots for up to ∼200 individuals with ∼ 40 chromosomes. For larger sampling sizes it is advised to generate a custom R code using the Clusters dataset generated by *pepa-base*. There are a total of three output plots, the “painted” chromosomes, and two barplots with the percentage of genome and gene content for each ancestry.

#### Quantifying Parental Contributions

Despite focusing on visualization, this tool includes a few quantitative measurements. In the pipeline *pepa-paint*, the code computes the percentage of the genome inherited from each parent for every individual by checking the size of the clusters with each ancestry. This code does not know the whole length of the genome and it uses only the length of the clusters. This is a simple process, however, it is surprisingly rare across tools for population genetics and provides a quantitative measure to accompany interpreting the painted chromosomes. Additionally, *pepa-paint* can calculate the percentage of genes inherited from each parent. These percentages can match, depending on the size of repeat regions and if biases in recombination exist. This gene-level analysis requires a GTF file, such as those available for many genomes in NCBI, or a user-generated annotation file. GTF files are available for many species, and the annotation format used can be easily made. By default, this code selects for all genes with the keyword ‘gene’ in a GTF file, however, this can be changed through a flag. By allowing users to define the types of genes included (e.g., protein-coding vs. non-coding RNAs), *pepa-paint* offers both simplicity and flexibility to perform a wide range of analyses.

## Results and discussion

To highlight the functionality of *pepa*, I analyzed the offspring of a cross between two strains of fission yeast (*Schizosaccaromyces pombe*). These strains have been identified before as belonging to two different ancestries, where the compatibility between strains and their phenotypes seems to be affected by the ancestry [14]. However, while work on identifying ancestry between strains has been performed before, we lack a tool to analyze the pedigree of an experiment, simply tracking from which parent they have inherited portions of their genome. We included here the results from the analyses made with *pepa* of whole genome sequencing of four offspring (**Fig.2**). The painted chromosomes (**Fig.2A**) clearly show that each offspring inherited most of Chr1 and Chr2 from the red strain, while Chr3 is mostly from the blue strain. These results align with previous work, where Chr3 has been found to host most of the wtf genes, meiotic drivers that kill offspring if not present in the genome. The painted Chr3 suggests that the blue strain has most of the wtf genes present in the red strain, but the red strain does not have the ones present in the blue one. Thus, the only offspring surviving are the ones inheriting enough Chr3. This inference matches our experience working with these strains, where crosses between these red and blue strains usually have only ∼5% surviving spores, as we obtained only four surviving offspring after culturing 72 individual spores. The inference from the painter chromosome suggests that recombination across regions seems random, but as there are only four samples, I cannot reliably test this effect. Nonetheless, the analysis shows that by using two clustering algorithms, *pepa* can assign and plot the ancestry of most of the genome across individuals.

**Figure 2.**
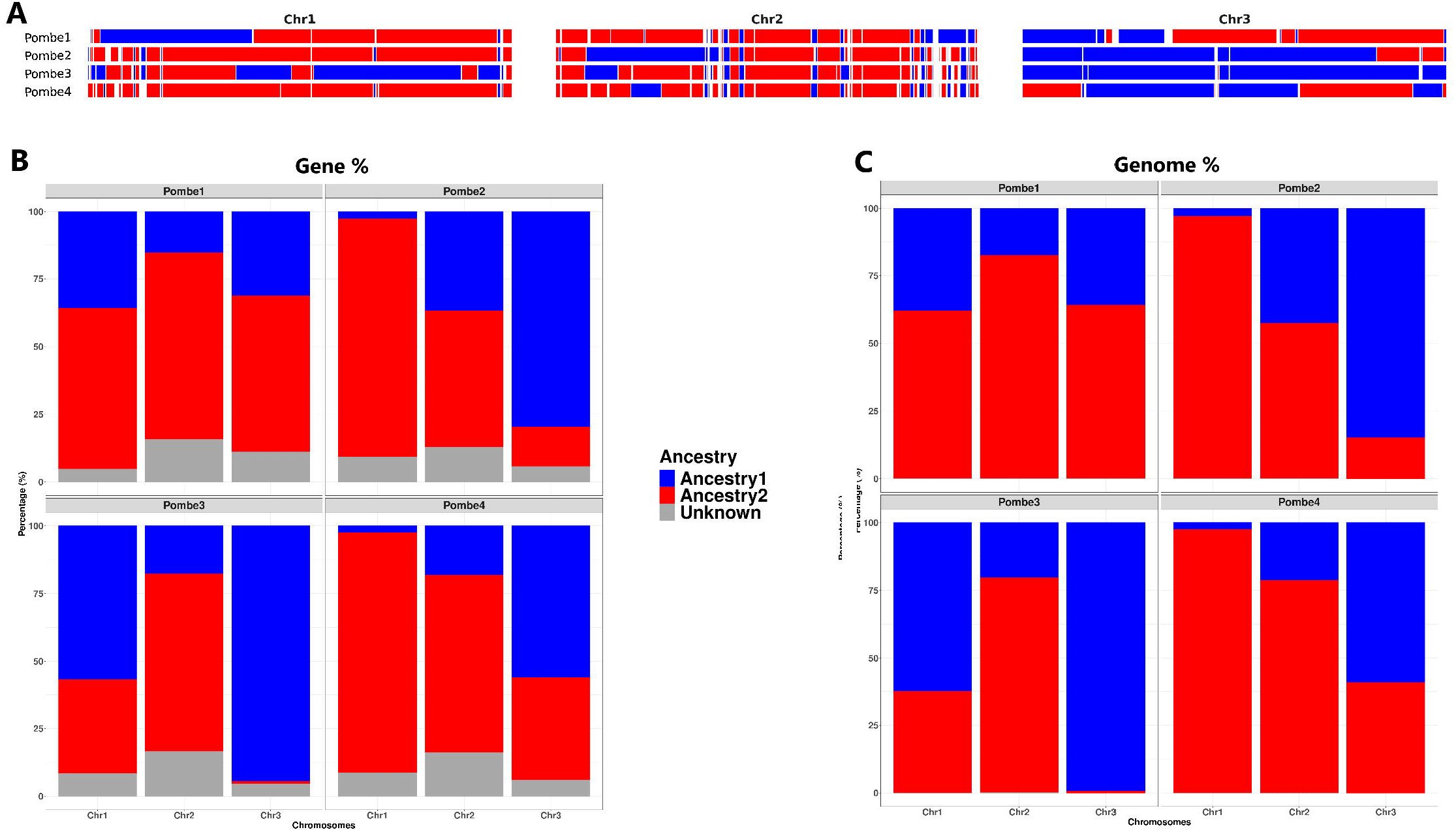
Graphical outputs of *pepa* analyzing samples from fission yeast. (A) Painted chromosomes for four individuals (Pombe1–Pombe4), showing inheritance from Ancestry1 (blue) and Ancestry2 (red). (B) Barplot showing the percentage of genes inherited from each ancestry across chromosomes (Gene %). Gray regions represent unknown ancestry, such as genes that belong to regions where the ancestry is not assigned. (C) Barplot showing the percentage of the genome inherited from each ancestry across chromosomes (Genome %). The ancestry in all plots is indicated always using the same colors for the same parents.

The quantitative measurements provided by *pepa* are similar, counting the percentage of protein-coding genes (**Fig.2B**) and nucleotides inherited from each parent (**Fig.2C**). The analysis shows that only the second offspring (Pombe2) has inherited ∼15% more Chr2 from the blue parent compared to the other offspring, however, the painted chromosome shows that this is due to a large recombination event in the first half of the chromosome. Similarly, Pombe1 and Pombe3 have much higher contributions from the blue parent, suggesting they should be similar strains, however, pairing this data with the painted chromosome contradicts this inference. The combined results show that the contribution of the blue parents for Pombe3 is on several regions not overlapping the regions in Pombe1, making these two strains as genetically different as the others. On the contrary, Pombe2 and Pombe4 seem more similar, with their main difference being a large recombination region on Chr3, a relatively small chromosome. This example highlights how the combination of different methods and data visualization can drastically affect the inference we can draw from the data.

## Conclusion

One of the key findings from using *pepa* is that genomic recombination does not occur uniformly across yeast chromosomes. Most of each chromosome tends to be inherited intact from one parent, with relatively few recombination events. This observation aligns with the expectation that recombination rates vary by genomic region and organism, but *pepa* makes these patterns easy to identify and quantify. The tool allows an intuitive visualization of recombination and inheritance patterns, and by limiting the dependencies necessary for installation, this tool is easily usable by both beginner and advanced users. Furthermore, the modular design ensures that it can be adapted to other organisms or extended for additional analyses, making it a valuable tool for exploring inheritance in diverse systems.

## Data availability

The tool can be found on GitHub and installed through conda (https://github.com/Mitopozzi/PePa) or by running directly the pipeline. The yeast data used for the analyses in VCF format can be found in the *Test* folder of the GitHub repository.

## Acknowledgments

I acknowledge the help of the student that help isolating the strains and extracting their DNA material, Luis Muxfeldt. Also, I want to thank Jochen Wolf for the insightful discussion and for convincing me to make a tool to paint chromosomes based on known ancestry. All writing, analyses, and interpretation were performed by AP.

